# Development of an immunoassay for the detection and diagnosis of microbially-influenced corrosion caused by methanogenic Archaea

**DOI:** 10.1101/2024.08.08.607177

**Authors:** Sven Lahme, Jorge Mantilla Aguas

## Abstract

Microbially-influenced corrosion (MIC) is a costly problem across several industries. The steadily rising use of advanced molecular biological methods to investigate MIC allowed ever deeper insights in the underlying microbial community structure and function. However, currently available technologies do not allow accurate detection and diagnosis of MIC in the field. Recent discovery of a special hydrogenase in certain corrosive methanogenic Archaea allowed for the development of a first ever MIC biomarker termed here *micH*. The *micH* gene encodes the large subunit of a special [NiFe] hydrogenase involved in MIC. Here we describe the development of a recombinant antibody that enable the specific and sensitive detection of the MicH protein in western blot immunoassays. Using a recombinant MicH protein we determined the lower limit of detection per assay to be around 0.3 fg MicH. The immunoassay was able to detect a strong signal for the MicH protein in *micH*-positive pure cultures of *Methanobacterium*-like. strain IM1 that was cultivated on iron granules, and the signal was over 500 times lower in a *micH*-negative *Methanococcus maripaludis* S2 culture. To further evaluate the ability to differentiate corrosive from non-corrosive microbial communities, we tested ten oil field enrichment cultures that showed a wide range of corrosion rates (0.02 – 0.48 mm/yr). We detected the MicH protein in planktonic (36.5 – 1473.5 pg/mL) and carbon steel biofilm samples (41.0 – 7971.3 pg/cm^2^) from corrosive methanogenic enrichments (0.17 – 0.48 mm/yr) but did not detect MicH in any of the non-corrosive tests (<0.08 mm/yr) despite methanogenic activity. The results indicate that corrosion was likely caused by methanogenic Archaea expressing a corrosive [NiFe] hydrogenase and that the newly developed MicH-specific immunoassay can detect and monitor their activity. The development of a specific and sensitive immunoassays to detect a MIC biomarker allow corrosion scientists and field practitioners to detect and monitor the activity of corrosive methanogenic Archaea.

## Introduction

Microbially-influenced corrosion (MIC) negatively affects steel infrastructures and causes significant costs across multiple industries ever year. In anaerobic systems, such as pipelines used for the production and transportation of mixtures of oil, water and gas, MIC is often linked to the presences and activity of sulfate-reducing bacteria (SRB) and methanogenic Archaea^1–5^.

The monitoring of corrosive biofilms in industrial settings often involves the quantification of cell numbers by culture-based or molecular biological methods such as the quantification of adenosine triphosphate (ATP) or universal and functional genes by quantitative PCR (qPCR)^6–9^. Despite the introduction of the latter into industrial operations, reliable detection of MIC during operation is still challenged. Firstly, not all microorganisms are causing industrially relevant corrosion and secondly, even members of the same metabolic groups (e.g. SRB or methanogenic Archaea) have been shown to have vastly different corrosion rates^1,10^. This suggest likely genetic differences that enable some microorganisms to cause higher rates of corrosion than others.

A recent study identified a specific gene islands in genomes of corrosive methanogenic Archaea that are absent in non-corrosive close relative strains. The gene islands encode among other genes a unique and phylogenetically highly conserved bidirectional [NiFe] hydrogenase (termed ‘MIC hydrogenase’), a Tat system for protein translocation as well as enzymes with potential involvement in protein glycosylation^5,11–13^. The ‘MIC hydrogenase’ was only detected in corrosive pure cultures and corrosive oil field enrichment cultures but not in the ones showing low corrosion, explaining the corrosive character of some methanogenic Archaea.

We recently proposed to use the large subunit of the ‘MIC hydrogenase’ as biomarker (termed *micH*) to detect and monitor certain corrosive methanogenic Archaea in the field^5^. By utilizing specific qPCR assays, the MIC biomarker showed excellent correlation with MIC in both laboratory tests and field applications^5,14^. Here, we discuss the development of a MicH-specific recombinant antibody to detect and monitor corrosive methanogenic Archaea in laboratory tests and in the field. In addition, protein detection may reduce potential false positive that could be associated with qPCR-based detection. A recombinant antibody able to detect MIC protein biomarker would also allow the development of a field-friendly rapid test assays (e.g. lateral flow tests) to support MIC monitoring and detection in the field.

## Materials and Methods

### Growth of pure cultures and enrichments

Actively growing cultures of *Methanococcus maripaludis* strain S2 (= DSM 14266), *Desulfovibrio ferrophilus* strain IS5 (= DSM 15579) and *Oleidesulfovibrio alaskensis* strain G20 (= DSM 17464) were acquired from the German strain collection DSMZ. Culture aliquots of *Methanobacterium*-like strain IM1 were donated by Dr Florin Musat (Helmholtz Centre for Environmental Research, Leipzig, Germany). Pure cultures and enrichments in this study were routinely grown in the presence of 1 – 2 mm iron granules (40 μg/ml medium; Thermo Fisher Scientific, Waltham, Massachusetts, US), 0.5 mM sodium acetate, 0.5 mM sodium sulfide and 10 mM sodium sulfate in anaerobic synthetic brine that was modelled after an oil field produced water as previously described^5^.

For establishing field enrichment cultures 100 ml Duran® glass bottles were filled with 2 g iron granules and dry heat sterilized at 180° C for 1 h. After cooling down the bottle headspace was exchanged with N2/CO2 (79:21) and sealed with sterile butyl rubber stoppers and screw cups. The sterile anoxic glass bottles were inoculated with 50 ml of produced water on site. Production fluids (oil, gas and water) were collected in sterile polypropylene bottles and oil and water allowed to separate by gravity. Once separated, the water was collected using sterile syringes with attached sterilized polypropylene tubes. After collection the tube was carefully replace by a sterile needle and the anoxic glass bottles inoculated. After arrival in the destination lab, the cultures were incubated at 32° C for at least two weeks before used as inoculum for corrosion bottle tests.

Enrichment cultures were established this way from eight pipelines of two separate West African oil field operations. In addition, two additional enrichment were established using pipeline solids (‘pigging debris’) collected as described previously^5^ from a pipeline from an U.S. West Coast operation and a U.S. onshore oil field operation during maintenance cleaning. Upon arrival in the lab approximately 3 g solids were transfer into sterile bottle containing 50 ml of the previously mentioned synthetic produced water^5^ and closed with sterile butyl rubber stoppers under N2/CO2 (79:21) atmosphere.

### Corrosion bottle tests

Corrosion tests were performed in one-liter Duran^®^ glass bottles that were filled with 600 ml anoxic synthetic brine to which three pre-cleaned and sterilized API 5L X52 steel coupons were added in polyether ether ketone coupon holder as described earlier^5,15^. The bottles were sealed with butyl rubber stoppers using sterile Hungate techniques and a N2/CO2 (79:21) headspace. Test were inoculated with 1% of respective cultures and incubated on a rotary shaker at 75 rpm and 32°C for eight weeks. Planktonic samples were collected for protein and DNA analysis using sterile syringes and 0.2 μm polyethersulfone filter membranes or sterile swabs for biofilm collection as mentioned previously^5^. Samples were immediately frozen at −80°C until further analysis. Removal of corrosion products and weigh loss corrosion analysis of corrosion coupons was performed at the end of the test as outlined earlier^5^.

### Recombinant antibody development

The recombinant antibody development workflow was conducted by Genscript Biotech (Piscataway, NJ) and will be briefly outline below. For the initial development of polyclonal antibodies, two peptide sequences were chosen from the protein sequence of the target gene *micH* of the corrosive methanogen *M. maripaludis* strain OS7 (see supplementary material for details) and conjugated to the metalloprotein keyhole limpet hemocyanin (KLH). Per peptide four rabbits were immunized with three additional consecutive boosters. After four months, blood was harvested from the rabbits and polyclonal antibodies partially purified for evaluation. Tests were performed by indirect enzyme-linked immunosorbent assay (ELISA) using either of the two peptides as target and via Western blotting as outlined below using protein extracts from *Methanobacterium*-like strain IM1 as positive control^16^ and protein extracts from *M. maripaludis* strain S2 was used as MicH-negative control (Fig. S1)^11^. Animals with best performing polyclonal antibodies were selected for cell fusion of spleen B cells with myeloma cell line. Screening and selection of clones were performed via immunoassays as described above. The best performing monoclonal antibody ‘A11-1’ was chosen for DNA sequencing (see supplementary material) and recombinant antibody production by Genscript Biotech.

### Protein extraction and detection via Western blot

Proteins were extracted from filters or swabs by placing them in sterile 2 ml screw cup microcentrifuge tubes. 900 μl lysis buffer (10 mM HEPES, 1% Triton X-100, 5 mM EDTA, 100 mM DTT and 100mM Na2CO3, 1% SDS, 12% sucrose) were added to the filters or swabs and placed in a heating block (105°C) for exactly 2 min. The tubes were removed and cooled to room temperature and then vigorously mixed and centrifuged for 5 min at 21,100 g. 200 μl of the supernatant was further processed to remove detergents using Pierce™ HiPPR Detergent Removal Spin Column Kit (Thermo Fisher Scientific, Waltham, MA) according to manufacturer’s protocol. The detergent-free protein samples were further processed using Pierce™ Protein Concentrators with a 10 kDa molecular weight cut-off polyether sulfone membrane (Thermo Fisher Scientific, Waltham, MA). Protein samples were concentrated to an approximate volume of 25 μl and either directly used for Wester blotting or stored at −80°C.

Western blotting was performed on a Protein Simple JESS Western blot platform (Protein Simple, San Jose, CA) using the 12 – 230 kDa Separation Module and the Jess Anti-Rabbit Detection Module. Protein analysis was performed according to the manufacturer protocol using a 1:20 antibody dilution (approx. 25 μg/ml) of A11-1 and 3 μl of protein sample. Protein Simple’s Total Protein module was used for normalization. For quantification a recombinant MicH protein was purchased from Genscript Biotech (Piscataway, NJ) developed following their large-scale *E. coli* expression protocol and His-tag purification.

### DNA extraction and qPCR analysis

For DNA extraction cells were collected via filtration and filters transferred to 5 ml cryo vials containing 1 ml DNA/RNA shield (Zymo Research, Irvine, CA) as preservative. Filters were stored at −80°C until further analysis. DNA extraction and *micH* qPCR analysis were performed by Microbial Insights Inc. (Knoxville, TN) as described previously^5^.

### Adenosine triphosphate analysis

ATP was quantified using the 2^nd^ Generation ATP^®^ test kit and PhotoMaster™ luminometer (LuminUltra Technologies Ltd., Fredericton, Canada). For analysis of planktonic cells 20 ml of medium was collected via filtration using the provided filters and syringes following the manufacturers protocol. Biofilms were collected using a minimum of two sterile swabs per coupon and the swab tips used for ATP quantification as outlined in the kit’s manual.

### Gas chromatography

Gas chromatographic analysis of methane was performed on an Agilent 490 Micro gas chromatograph using a thermal conductivity detector. Separation was achieved on a heated Agilent PoraPlot U column (50°C, 22.1 lb/in^2^, and 10 m) in argon gas.

## Results

### Recombinant Anti-MicH antibody showed excellent affinity for recombinant MicH protein

In order to optimize the signal to noise ratio (S/N) we performed first a dilution series with the Anti-MicH antibody and a fixed concentration of 1 ng/ml of MicH_rec_. A signal was detected between approximately 50 – 60 kDa molecular weight that is in good agreement with the expected molecular weight of the MicH protein containing the His-tag used for purification (54.5 kDa; Fig. 1A). Test performed with different antibody dilutions of Anti-MicH showed best results for S/N ratio and low baseline with dilutions 1:50, 1:20 and 1:10 (Fig. 1B). For subsequent tests an antibody dilution of 1:20 was chosen to minimize potential background noise while achieving an overall strong signal.

**Figure 1.**
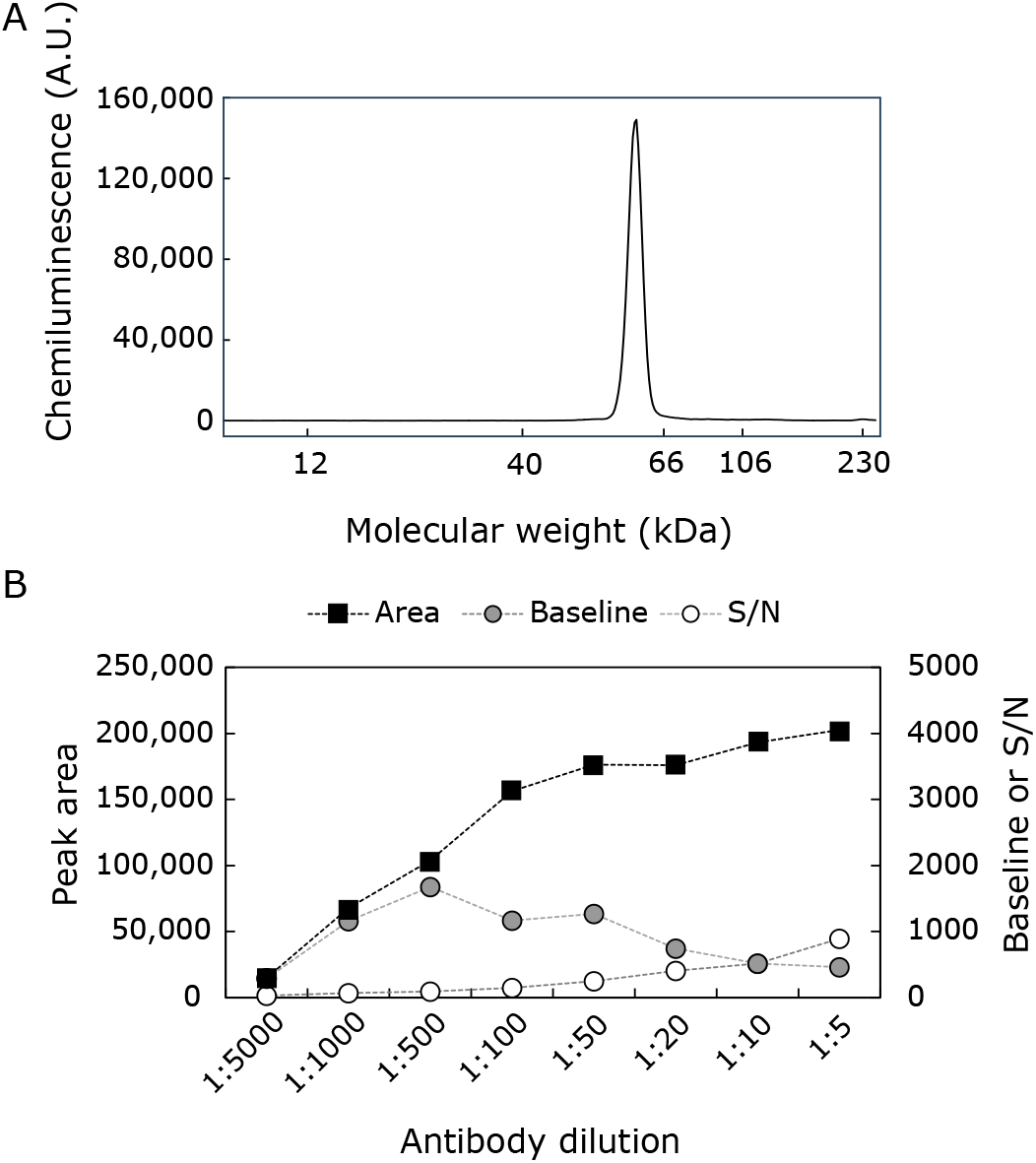
Performance of recombinant antibody Anti-MicH using a recombinant MicH protein as target. A) Chemiluminescence measured as arbitrary units (A.U.) showing signal at the expected molecular weight of MicH_rec_. Test was performed with a 1:20 dilution of the antibody. B) Results of tests performed with varying dilutions of Anti-MicH and 1 ng/ml MicH_rec_.

In addition, a calibration curve of recombinant MicH protein showed excellent linear response with concentrations between 20 pg/ml and 10 ng/ml (Fig. 2 and Fig. S2). A clearly detectable signal with acceptable S/N ration (>30) was still detected at 20 pg/ml MicH_rec_ therefore the assay was able to reliably detect as little as 0.3 fg of recombinant MicH corresponding to approximately 3·10^3^ protein molecules.

**Figure 2.**
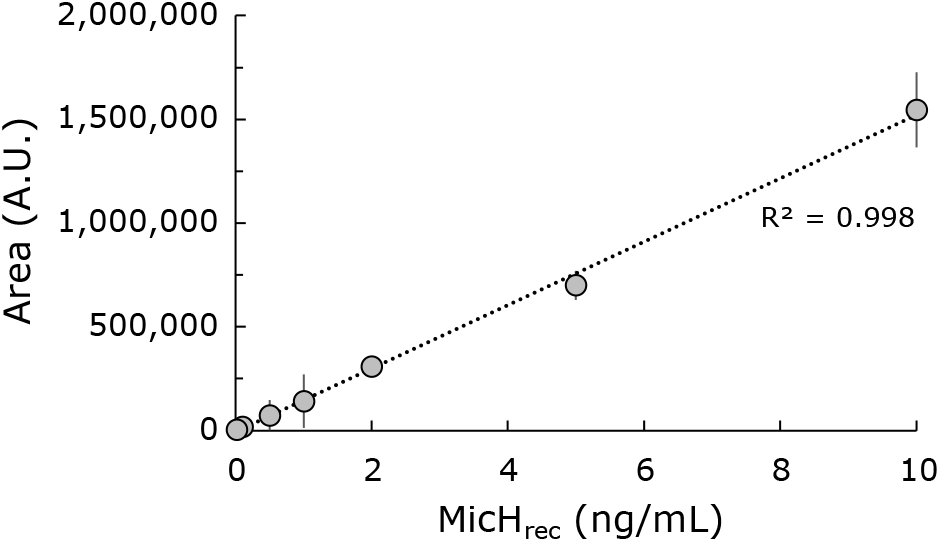
Western Blots performed with dilution series of recombinant MicH protein and Anti-MicH antibody dilutions of 1:20. The error bar represent the standard deviation of technical triplicates.

### MicH-specific immunoassay showed excellent selectivity for corrosive methanogenic cultures

Unlike qPCR assays, immunoassays are less selective and may show potential background signal due to cross-reactivity with non-target proteins or peptides in absence or low concentration of their specific target^17^. To test for assay specificity and the ability to distinguish between corrosive and non-corrosive methanogenic Archaea as well as non-methanogenic cultures, we tested the MicH-specific immunoassay on protein extracts from methanogenic and sulfate-reducing pure cultures (Fig. 3).

**Figure 3.**
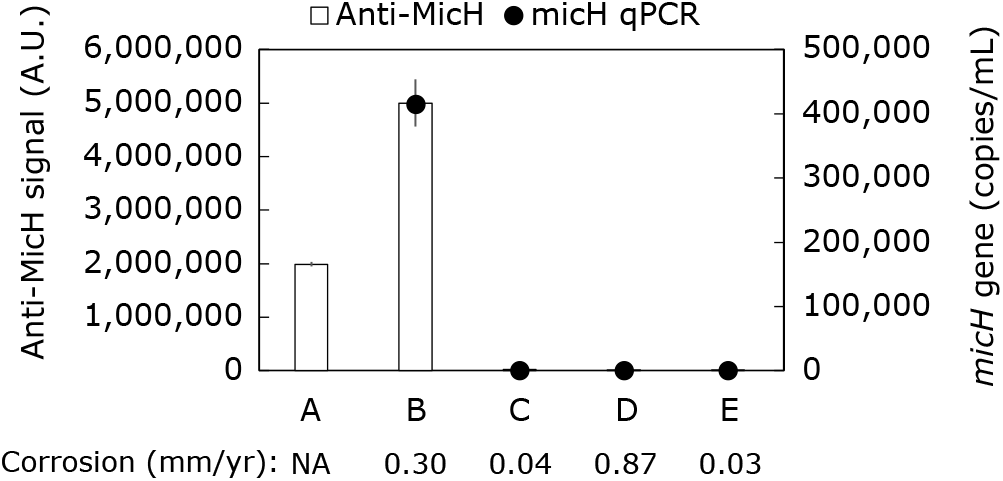
Analysis of protein extracts with MicH-specific Western blot immunoassay and DNA extracts with micH qPCR assay. A) Recombinant MicH Protein (10 ng/mL), B) *Methanobacterium* sp. strain IM1, C) *Methanococcus maripaludis* strain S2, D) *Desulfovibrio ferrophilus* IS5, E) *Desulfovibrio alaskensis* G20. Average corrosion rates are shown below the bar plot. The error bars where seen indicate the standard deviations of triplicate cultures or technical triplicates in case of recombinant MicH.

Protein extract of *Methanobacterium*-like strain IM1 obtained from cultures, that inflicted significant corrosion of 0.3 mm/yr, showed a strong signal in the Western blot (Fig. 3 and Fig. S3A). Furthermore, the *micH* gene was also detected by qPCR in DNA extracted from cultures of strain IM1, confirming both the presence and expression of the ‘MIC hydrogenase’ during corrosion. In contrast, the MicH-specific assay showed a more than 500x lower signals for protein extracts from *M. maripaludis* strain S2 known to lack the ‘MIC hydrogenase’^11^. In agreement, strain S2 caused only negligible corrosion of 0.04 mm/yr (Fig. 3). Similar low signals were observed in protein extracts from *Desulfovibrio ferrophilus* IS5 and *Desulfovibrio alaskensis* G20. DNA analysis via qPCR confirmed absence of the *micH* gene in those cultures (Fig. 3). The results confirm that the MicH-specific immunoassay selectively detects the MicH protein while showing low cross-reactivity with proteins extracted from non-corrosive methanogenic strain S2 and SRB pure cultures (Fig. 3 and Fig. S3A).

### MicH-specific immunoassay allowed detection of MicH in corrosive methanogenic field cultures

Next, we wanted to better understand the performance of the MicH-specific immunoassay and selectivity of the Anti-MicH antibody in more complex context of oil field produced water enrichment cultures. Therefore, ten field enrichment cultures that showed varying degree of MIC activity were compared in terms of corrosion as well as presence of the *micH* gene and expression of the MicH protein (Fig. 4). Five of the ten cultures were classified as ‘+MIC’ showing averaged corrosion rates between 0.48 – 0.17 mm/yr, which according to the Association for Materials Protection and Performance (AMPP) is considered high to severe (AMPP-SP0775-2013). The other five cultures were classified as ‘-MIC’ showing rates between 0.02 – 0.08 mm/yr, considered as low to moderate corrosion according to AMPP (Fig. 4A).

**Figure 4.**
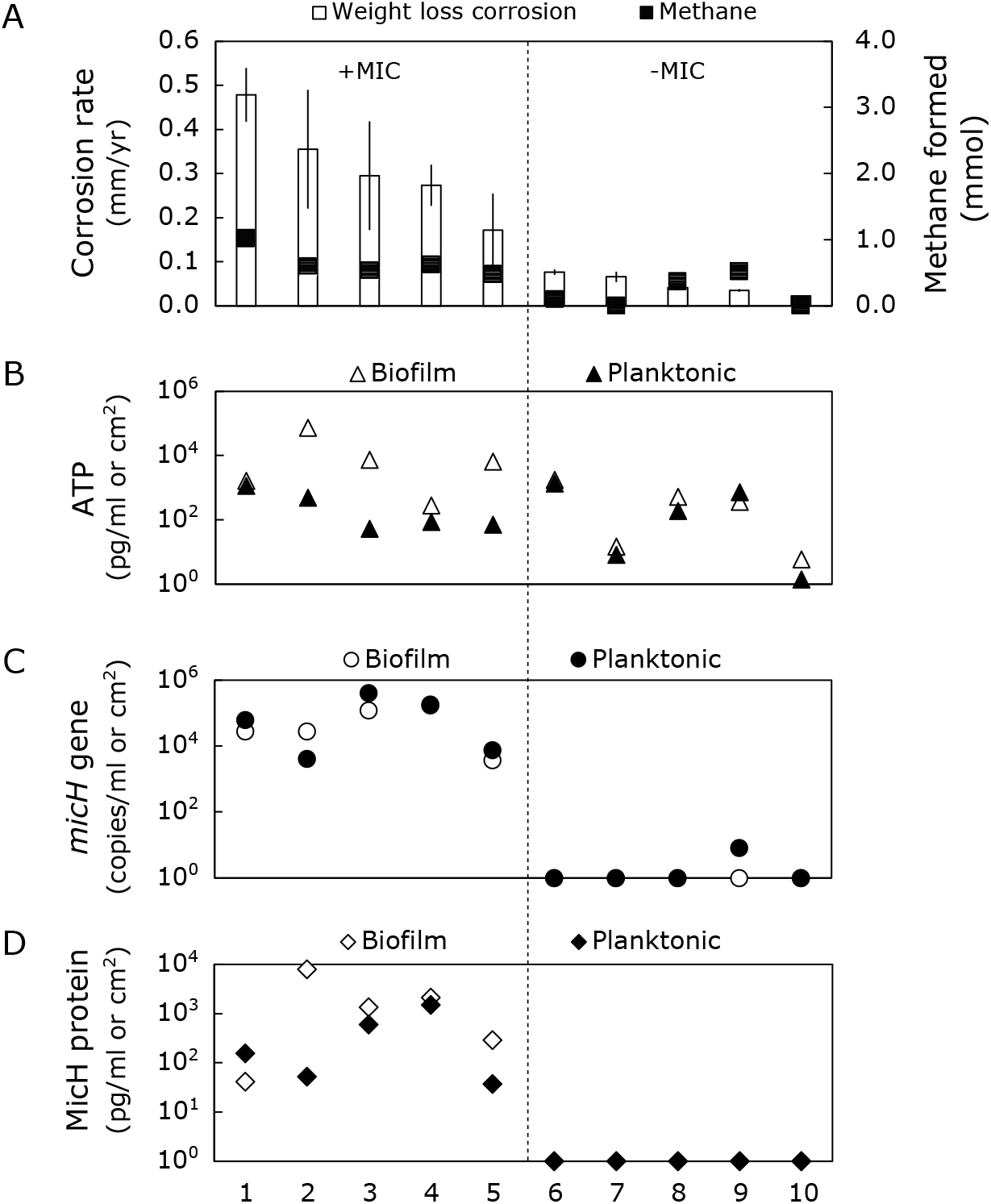
A) Weight loss corrosion rates and methane formation in oil field produced water enrichment cultures classified as corrosive [‘+MIC’] and non-corrosive [‘-MIC’] enrichments. B) MicH protein numbers detected in corrosion coupon biofilms (open diamonds) and planktonic cells (closed diamonds). Enrichments 1, 2, 4, 6, 7, 9, 10 – First West African oil field. Enrichment 3 – Second West African oil field. Enrichment 5 – U.S. offshore oil field. Enrichment 8 – U.S. onshore oil field. Error bars in A) indicate standard deviation of weight loss analysis from three corrosion coupons.

Of the ten cultures eight showed detectable methanogenic activity having formed between 0.1 and 1.0 mmol of methane at the end of the test (Fig. 4A). ATP results indicated that biofilms have formed on steel surfaces to a varying degree with generally higher numbers detected on corroded steel surfaces (+MIC; 278 – 71299 pg/cm^2^) compared to less corroded ones (-MIC; 6 – 1762 pg/cm^2^; Fig. 4B). The *micH* gene was detected in planktonic cells as well as all five biofilms collected from corroded steel coupons (>0.17 mm/yr; Fig. 4C). The MicH protein was similarely detected in planktonic (36.5 – 1473.5 pg/mL) as well as biofilm samples (41.0 – 7971.3 pg /cm^2^) collected from ‘+MIC’ enrichments, indicating active expression of the ‘MIC hydrogenase’ in those cultures (Fig. 4D and Fig. S3B). Interestingly, we detected a small amount of the *micH* gene in one of the enrichments showing low corrosion (0.04 mm/yr). However, the MicH protein has not been detected, suggesting that the *micH* gene may not have been expressed and no corrosive biofilm formed (Fig. 4).

## Discussion

Multiple studies have highlighted the importance of understanding the underlying MIC mechanisms to better detect and diagnose MIC^5,7,18,19^. Here we presented the development of a MicH-specific immunoassay that showed strong affinity for the large subunit of a the ‘MIC hydrogenase’ shown to be involved in corrosion by certain methanogenic Archaea^5,11^. The results obtained with pure and field enrichment cultures suggest that the Anti-MicH antibody is highly selective for the MicH protein found in certain corrosive methanogenic Archaea while showing little to no cross-reactivity with proteins from SRB or non-corrosive methanogenic Archaea. This also included oil field enrichment cultures showing methanogenic activity. In addition, the assay confirmed that corrosive methanogenic cultures indeed actively expressed the ‘MIC hydrogenase’ thereby accelerating the iron oxidation of carbon steel as previously shown in pure cultures and genetically modified *Methanococcus maripaludis* strains^11,12^.

Interestingly, we were able to detect the *micH* gene as well as the MicH protein in cultures of the corrosive *Methanobacterium*-like strain IM1^16^. To our knowledge this is the first dataset that allowed a deeper insight into the underlying molecular mechanisms of this corrosive methanogenic archaeon. Previous potential step measurements have suggested a biologically catalyzed H2 production pathway on graphite electrodes^20^. The identification of the *micH* gene and protein in strain IM1 further supports an enzymatic catalyzed pathway for proton reduction on steel surfaces.

The detecting of the MicH protein not only in biofilms but also planktonic cells suggest that the assay may be applicable to both MIC monitoring in biofilms as well as water samples. The latter is often readily available in the field, while it is often challenging to obtain relevant biofilm samples in the field to monitor pipeline corrosion. Biofilms act as planktonic cell factories continuously shedding cells into fluids streams^21,22^ that can then be collected at pipeline outlets and can be analyzed for presents and expression of the MIC biomarker. We could previously show the feasibility of using qPCR assays to detect MIC biomarkers in produced water samples as well as pipeline solids from various global operations ^5,14^ and it would be interesting to see whether similar can be accomplished using the novel immunoassay.

Overall, the development of a MicH-specific immunoassay offers an additional tool for corrosion scientists and field practitioners to detect, monitor and understand corrosive methanogenic biofilms. Furthermore, the detection of the MicH protein in multiple field enrichment cultures from different locations together with previous qPCR results^5^ highlight the importance of this MIC mechanism in anaerobic systems.

## Conclusion

Here, we built on previous research and findings to develop specific immunoassay for the detection of an industrially important MIC mechanism. The assay offers corrosion scientists and field practitioners a novel way to detect, monitor and understand corrosive methanogenic biofilms. Furthermore, the detection of the MicH protein in multiple field enrichment cultures from different locations together with previous qPCR results5 highlight the importance of this MIC mechanism in anaerobic systems on a global scale.

□ Establishment and optimization of a novel Western blot-based immunoassay to detect an important MIC biomarker (‘MicH’) found in certain highly corrosive methanogenic Archaea.
□ Analysis of recombinant MicH protein and pure cultures of corrosive and non-corrosive methanogens and SRB showed high sensitivity and selectivity of the immunoassay
□ Test with corrosive and non-corrosive oil field enrichment cultures showed applicability for Mic detection in the field.

## Acknowledgement

The authors like to thank Manuel Castelan for assistance with gas analysis and Jaspreet Mand for providing protein extracts for the study as well as Dennis Enning for the fruitful discussions. In addition, we are grateful to Florin Musat for providing a culture of strain IM1 for our study.

