## Supplementary material for "Development of an immunoassay for the detection and diagnosis of microbially-influenced corrosion caused by methanogenic Archaea"

Protein sequence of the large subunit of the ‘MIC hydrogenase’ (referred to as MicH*)* from *Methanococcus maripaludis* strain OS7. The peptide sequence used for antibody development is highlighted in yellow.

>WP_119846095.1

MAVEIKPVTRIEGDGKLELETYLVDGKLRVKNLLTPTDGTLNLPTDNAARYPKFCVTEFRGFEKF
AVGEQPETVTKLVSRICGVCPVPQNMASTCAVEAAYGTNIVDNAKAVRRLMLAVHTVHSHLLHF
FVLAGKDMLPHSIIDQELPNIISAHNKAQACVAVFGGKPVHPASCIPGGQTKVPNATEIGQIKARM
QEIQSYVVSLLGTLKGALLALPNDFGIRPCNYMSSGVPFYSGVSGSFSYFGTSGGVVIRDPSNPTD
FNSATIAPFNPENVKEEVITPYGSVISLPTNYSYTKRPYYAYNGVNYVCEVGPLARLGVAYKAGDS
VVKSTVDSIASDWGVAANDILSPSTRNRHITRLIETVIMIEMILNGGWSNIAHVDSYSPPNEKSGY
GVGVIEAPRGTLIHRIRVGSDLKTASYDCMVPTTANTGAFEEAVADDLVEIRPETISLLGRDPTEDE
KLLLGDASRTIRGFDPCCSCSSHMIEIKNPDGTIDGTK

DNA sequence of the heavy and light chains of the recombinant Anti-MicH antibody ‘A11-1’

>A11_1_Heavy_chain

ATGGAGACTGGGCTGCGCTGGCTTCTCCTGGTCGCTGTGCTCAAAGGTGTCCAGTGTCAGTC
GGTGGAGGAGTCCGGGGGTCGCCTGGTCACGCCTGGGACACCCCTGACACTCACCTGCACA
GTCTCTGGATTCTCCCTCAGTAGGTATGCAGTGAGCTGGGTCCGCCAGGCTCCAGGGGAGGG
GCTGGAATGGATCGGAGTCATTAGTGCTGGTGATAACACATACTACGCGAGCTGGGCGAAAG
GCCGATTCACCATCTCCAAAACCTCGTCGACCACGGTGGATCTGAAAATCACCAGTCCGACA
ACCGATGACACGGCCACCTATTTCTGTGCCAGAGGACCTTATGCTAATGATAACATCTGGGGC
CCAGGCACCCTGGTCACCGTCTCCTCA

>A11_1_light_chain

ATGGACACGAGGGCCCCCACTCAGCTGCTGGGGCTCCTGCTGCTCTGGCTCCCAGGTGCCAC
ATTTGCCGCCGTGTTGACCCAGACTCCATCTCCCGTGTCTGCAGCTGTGGGAGGCACAGTCAG

CATCAGTTGCCAGTCCACTGGTCGTCTTGTTAATAATAATTGGTTATCCTGGTTTCAGCAGAAA
CCAGGGCAGCGTCCCAAGCTCCTGATCTATCTGGCGTCCACTCTGGCATCTGGGGTCCCATCG
CGGTTCAGCGGCCGTGGATCTGGGACACAGTTCACTCTCACCATCAGCGACGTGCAGTGTGA
CGATGCTGCCACTTATTATTGTGTGGGCGGTAGTGATAGTGGTAGTAGTGATAATGGTTTCGGC
GGAGGGACCGAGGTGGTGGTCAAA

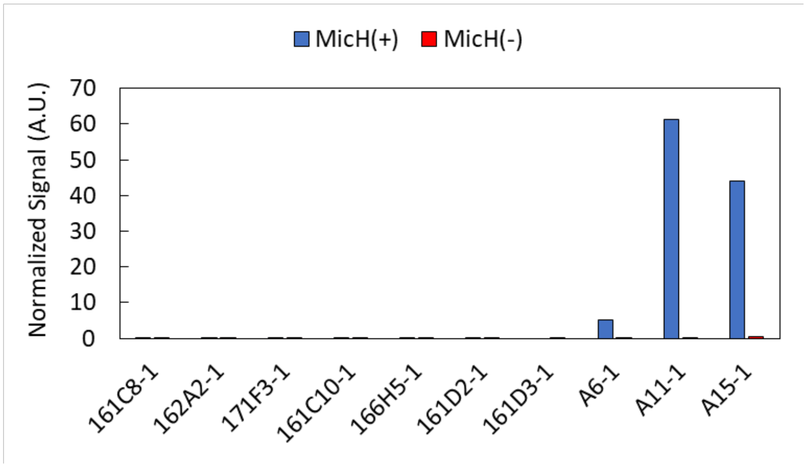


Figure S1 Selection of best performing Anti-MicH antibody using automated capillary Western Blotting. Recombinant monoclonal antibody A11-1 showed strongest signal with MicH(+) *Methanobacterium*-like strain IM1 and low cross reactivity with MicH(-) *Methanococcus maripaludis* strain S2.

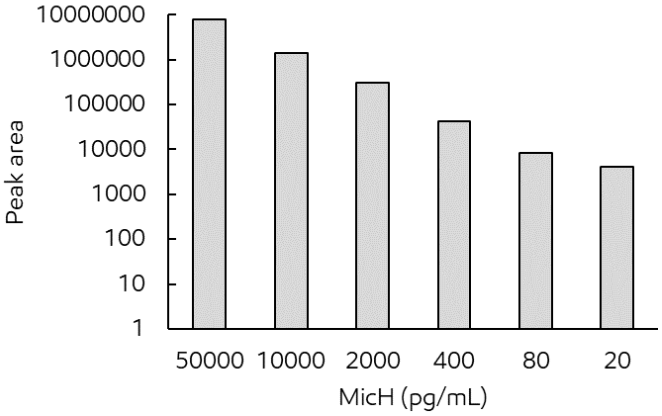


Figure S2 Western blot calibration curve of recombinant MicH protein between 50 ng/ml and 20 pg/ml using an Anti-MicH antibody dilution of 1:20.

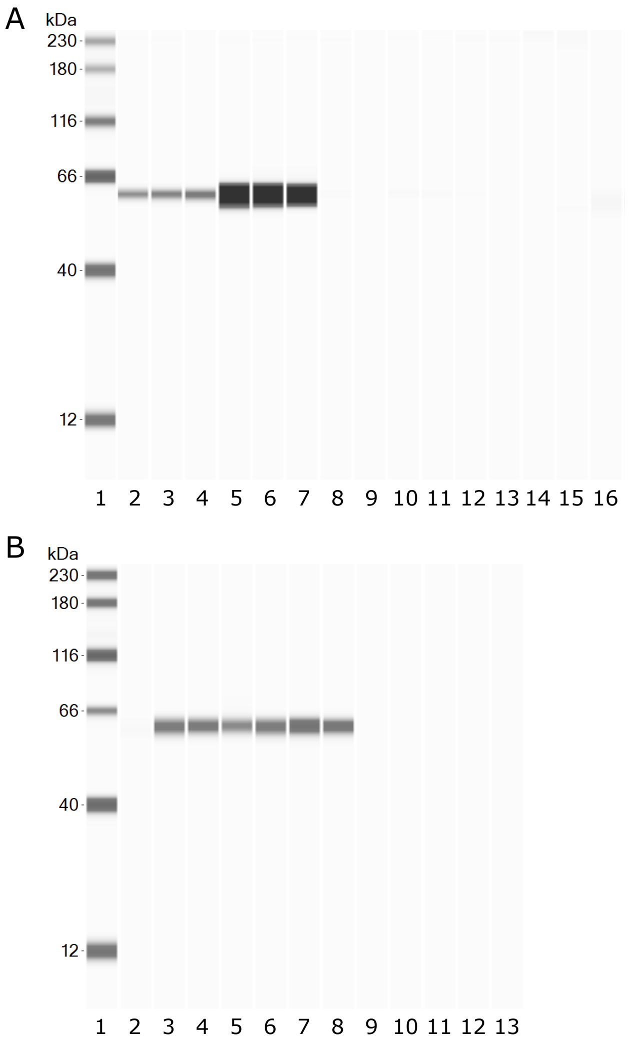


Figure S3 Computer-generated gel images of capillary Western blot runs of protein extracts of triplicate pure cultures grown in the presence of iron coupons. Lane 1: Protein ladder; Lanes 2-3: recombinant MicH (10 ng/ml); Lanes 5-7: *Methanobacterium*-like strain IM1; Lane 8-10: *Methanococcus maripaludis* strain S2; Lane 11-13: *Desulfovibrio ferrophilus* strain IS5; Lane 14-16: *Desulfovibrio alaskensis* strain G20. B) Analysis of protein extract from field enrichment cultures (planktonic samples) at the end of the corrosion bottle tests. Lane 1: Protein ladder; Lane 2: negative control; Lane 3: recombinant MicH (10 ng/ml); Lane 4: Enrichment 1; Lane 5: Enrichment 2; Lane 6: Enrichment 3; Lane 7: Enrichment 4; Lane 8:Enrichment 5; Lane 9: Enrichment 6; Lane 10: Enrichment 7; Lane 11: Enrichment 8; Lane 12: Enrichment 9; Lane 13: Enrichment 10.
